# Aging in zebrafish is associated with reduced locomotor activity and strain-dependent changes in predator avoidance behaviors related to anxiety

**DOI:** 10.1101/2023.11.03.565457

**Authors:** Jacob Hudock, Justin W. Kenney

**Affiliations:** Department of Biological Sciences, Wayne State University, Detroit, MI 48202

## Abstract

Aging is associated with a wide range of physiological and behavioral changes in many species. Like humans, zebrafish exhibit gradual senescence, and thus may be a useful model organism for identifying evolutionarily conserved mechanisms related to aging. Here, we compared behavior in the novel tank test of young (6-month-old) and middle aged (12-month-old) zebrafish from two strains (TL and TU) and both sexes. We find that this modest age difference results in a reduction in locomotor activity and strain dependent changes in predator avoidance behaviors related to anxiety. Older TL fish have an elevation in bottom dwelling whereas older TU fish have a decrease in thigmotaxis. We found no consistent effects of age on either short-term (within session) or long-term (1 day later) habituation to the novel tank. Our findings support the use of zebrafish for the study of how age affects locomotion and how genetics interacts with age to alter the regulation of emotional behaviors in response to novelty.

## Introduction

The challenge of understanding how the detrimental effects of age can be mitigated is of central concern in medicine and biology and is facilitated by the comparative study of aging across species (Arking, 2006; Austad, 2009). Zebrafish, like humans, display gradual senescence (Kishi et al., 2009). With advancing age, zebrafish have reduced regenerative capacity, accumulation of DNA damage, and cardiovascular impairment (Gilbert et al., 2014; Sun et al., 2014). The shared consequences of aging in humans and zebrafish, combined with the low cost of housing zebrafish for extended periods, suggests that zebrafish could serve as a valuable model organism in the study of aging.

In humans, many behaviors vary across the lifespan, such as emotional regulation and anxiety (John and Gross, 2004; Lenze and Wetherell, 2011; Nivard et al., 2015). The trajectory of these behaviors is shaped by the complex interplay of environmental factors with genetics and sex (Nivard et al., 2015; Singh et al., 2019). Untangling the web of interactions between age, genetics, and sex, is facilitated by use of model organisms for which many molecular genetic tools are available, like zebrafish. Although there has been a large increase in the study of zebrafish behavior in the past twenty years (Gerlai, 2023; Kenney, 2020), only a handful of reports have explored age-associated behavioral alterations. These studies have found that older fish have changes in circadian rhythms, decreased associative learning, and reduced locomotion (Gilbert et al., 2014; Philpott et al., 2012; Ruhl et al., 2015; Yu et al., 2006; Zhdanova et al., 2008). However, it is unknown if aging also affects commonly studied behaviors related to emotional regulation and exploration in zebrafish.

One of the most widely used behavioral tasks to study exploratory and emotional behavior in zebrafish is the novel tank test (NTT). In this test, animals are placed into a tank with which they have no prior experience. Spending time near the bottom or periphery of the tank are typically interpreted as fearful predator avoidance behaviors related to anxiety (Champagne et al., 2010; Gerlai et al., 2000; Kalueff et al., 2013; Levin et al., 2007; Maximino et al., 2010). Nonassociative memory can also be assessed by measuring how behavior habituates within session (short-term memory) or between daily sessions (long-term memory). To determine whether age affects behavior in the NTT, and if the effects of age are modulated by sex and genetics, we examined behavior in young (6 months old) and middle aged (12 months old) fish from two strains (TL and TU) and both sexes. We find that this modest difference in age is sufficient to alter several behaviors such as locomotion and predator avoidance.

## Results

### The effects of age on predator avoidance behaviors

We measured the behavior during exploration of a novel tank in young (6 mpf) and old (12 mpf) zebrafish from both sexes and two strains, TLs and TUs. Distance from the bottom of the tank is considered a predator avoidance response that is one of the most widely studied behaviors in zebrafish (Gerlai, 2010; Kalueff et al., 2013). This behavior has also been interpreted as ‘anxiety-like’ where spending more time near the bottom of the tank indicates greater anxiety (Maximino et al., 2010). In TL fish, there was a medium effect of age on distance from the bottom of the tank (Figure 1A, top; P = 0.035, η^2^ = 0.08) where older fish had greater anxiety-like behavior, spending more time near the bottom of the tank. There was no effect of sex (P = 0.36) or an interaction (P = 0.21). In TU fish, age did not affect bottom dwelling (Figure 1B, bottom, P = 0.14) and there was no effect of sex (P = 0.23) or an interaction (P = 0.30).

**Figure 1.**
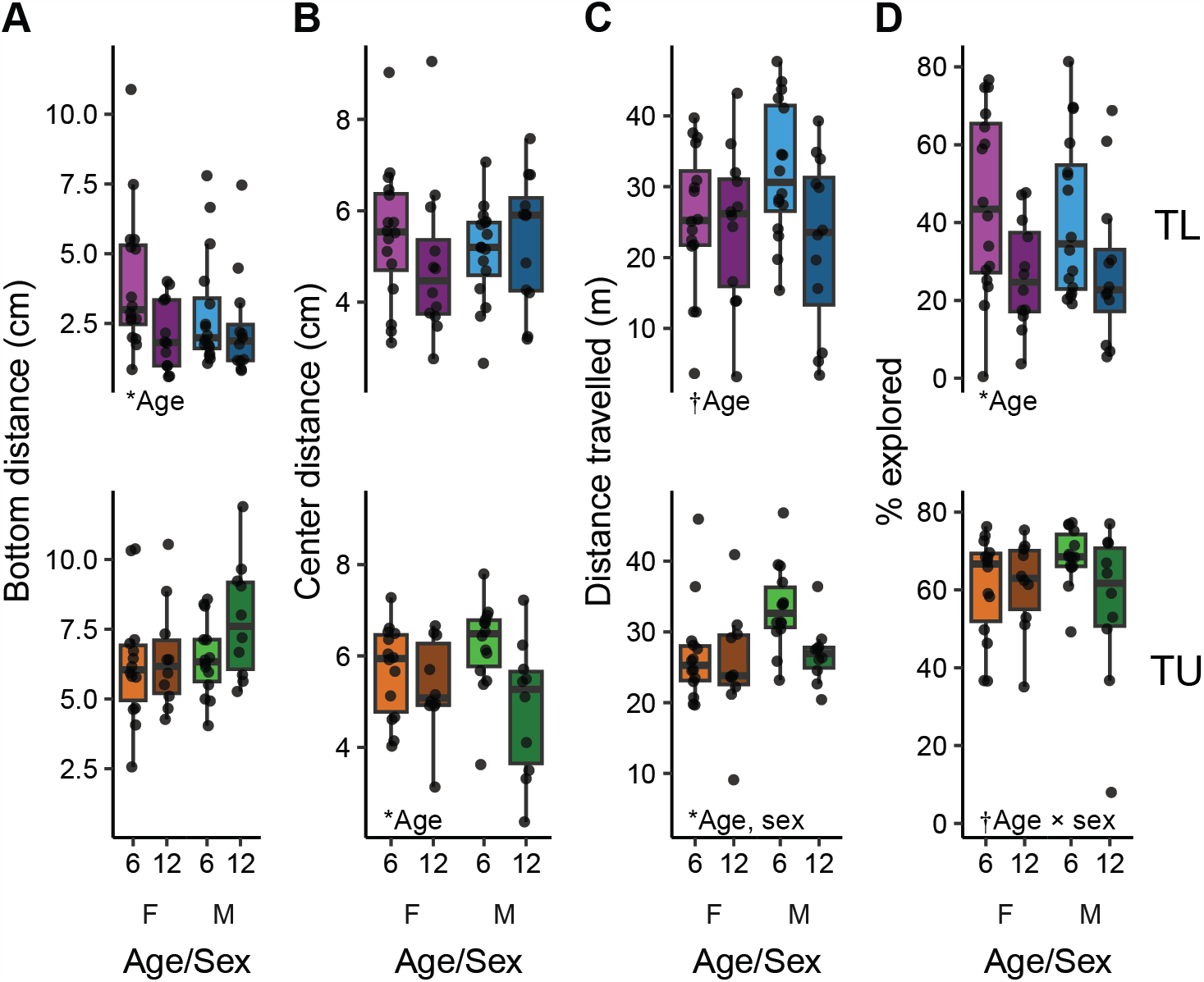
Overall behavior during initial exposure to the novel tank in young (6 mpf) and old (12 mpf) TL (top) and TU (bottom) fish of both sexes. Behaviors assessed were (A) bottom distance, (B) center distance, (C) distance travelled and (D) percent of the tank explored. Boxplot center line is the median, hinges are the interquartile range, and whiskers are hinges ± 1.5 times the interquartile range. *P < 0.05, †P < 0.10 from 2 × 2 (age × sex) ANOVAs.

Thigmotaxis (i.e. time spent near the periphery) of a tank is also commonly interpreted as a predator avoidance response in zebrafish, although this interpretation has recently come under question (Borba et al., 2023; Rajput et al., 2022). In TL fish (Figure 1B, top), there were no effects of age (P = 0.68), sex (P = 0.96) or an interaction (P = 0.23) on center distance.

However, in TU fish (Figure 1B, bottom), there was a medium-sized effect of age (P = 0.013, η^2^ = 0.12) where older fish spent more time in the center of the tank than younger fish. There was no effect of sex (P = 0.92) or an interaction (P = 0.12). Taken together with the findings on bottom distance, the effects of age on predator avoidance/anxiety-like behaviors depends on the strain of the fish: older TL fish spend more time near the bottom of the tank whereas older TU fish spend more time in the center of the tank.

### The effects of age on locomotion and exploration

Aging in zebrafish has been associated with a decrease in activity levels (Philpott et al., 2012). To determine if this also occurred during the NTT, we measured how far fish travelled during their exposure to the novel tank (Figure 1C). In TL fish, there was a strong trend towards a medium-sized effect of older fish swimming less than younger fish (P = 0.057, η^2^ = 0.064) with no effect of sex (P = 0.32) or an interaction (P = 0.12). In TU fish, there was a medium-sized effect of age (P = 0.044, η^2^ = 0.077) with older fish also swimming less than younger fish. There was also a slightly larger effect of sex in TUs (P = 0.022, η^2^ = 0.10) where males swam further than females; there was no interaction between age and sex (P = 0.17). Thus, older animals swim less than younger animals, an effect that appears to be largely independent of strain.

To determine if age influences how much of the novel tank fish choose to explore, we measured the percentage of the tank visited by fish (Figure 1D). In TL fish, there was a strong effect of older fish exploring less of the tank than younger fish (P = 0.0052, η^2^ = 0.14). There was no effect of sex (P = 0.79) or an interaction (P = 0.61). In TU fish, there was no effect of age (P = 0.14) or sex (P = 0.54) on percent explored. There was a trend towards a small interaction effect (P = 0.10, η^2^ = 0.055), although follow-up post-hoc tests within sex did not uncover any age differences (female: P = 0.91, male: P = 0.18). These findings suggest that background strain strongly modulates the influence of age on the willingness to explore a new environment, having a bigger impact on TLs than TUs. This is unlikely to be due entirely to less locomotor activity in older fish because older TU fish swam less than their younger counterparts with no impact on percent explored.

### The effects of age on short-term habituation

When exposed to a novel environment many animals, including fish, habituate such that their behaviors change within session (Levin et al., 2007; Rankin et al., 2009; Wong et al., 2010). To determine if habituation occurred during exposure to the novel tank, and whether this habituation was affected by age, we looked at how predator avoidance behaviors and locomotion changed between the first two and last two minutes in the novel tank (Figure 2). For changes in bottom distance in TL fish (Figure 2B, top), we found no effects of age (P = 0.40), sex (P = 0.66) or an interaction (P = 0.86). When compared to zero (i.e., no change), we did find strong effect in older male animals, which increased their bottom distance over time (P = 0.0094, d = 1.1), while younger males and older females had trends towards medium sized increases in bottom distance (P’s = 0.063 & 0.065, d’s = 0.59 & 0.64, respectively) and young females did not change (P = 0.16). The lack of effect in young TL females may reflect the fact that they had high bottom distance from the beginning of the trial (Figure 2A). In TU fish there were medium size effects of age (P = 0.030, η^2^ = 0.094) and sex (P = 0.045, η^2^ = 0.079) with no interaction (P = 0.59): older fish increased bottom dwelling over time (Figure 2B, bottom). When comparing the difference to zero, however, the results were less clear. Older female fish had a trend towards a strong effect of decreasing bottom distance (P = 0.066, d = 0.93) whereas young females (P = 0.84), young males (P = 0.27), and older males (P = 0.84) did not change their bottom dwelling behavior over time.

**Figure 2.**
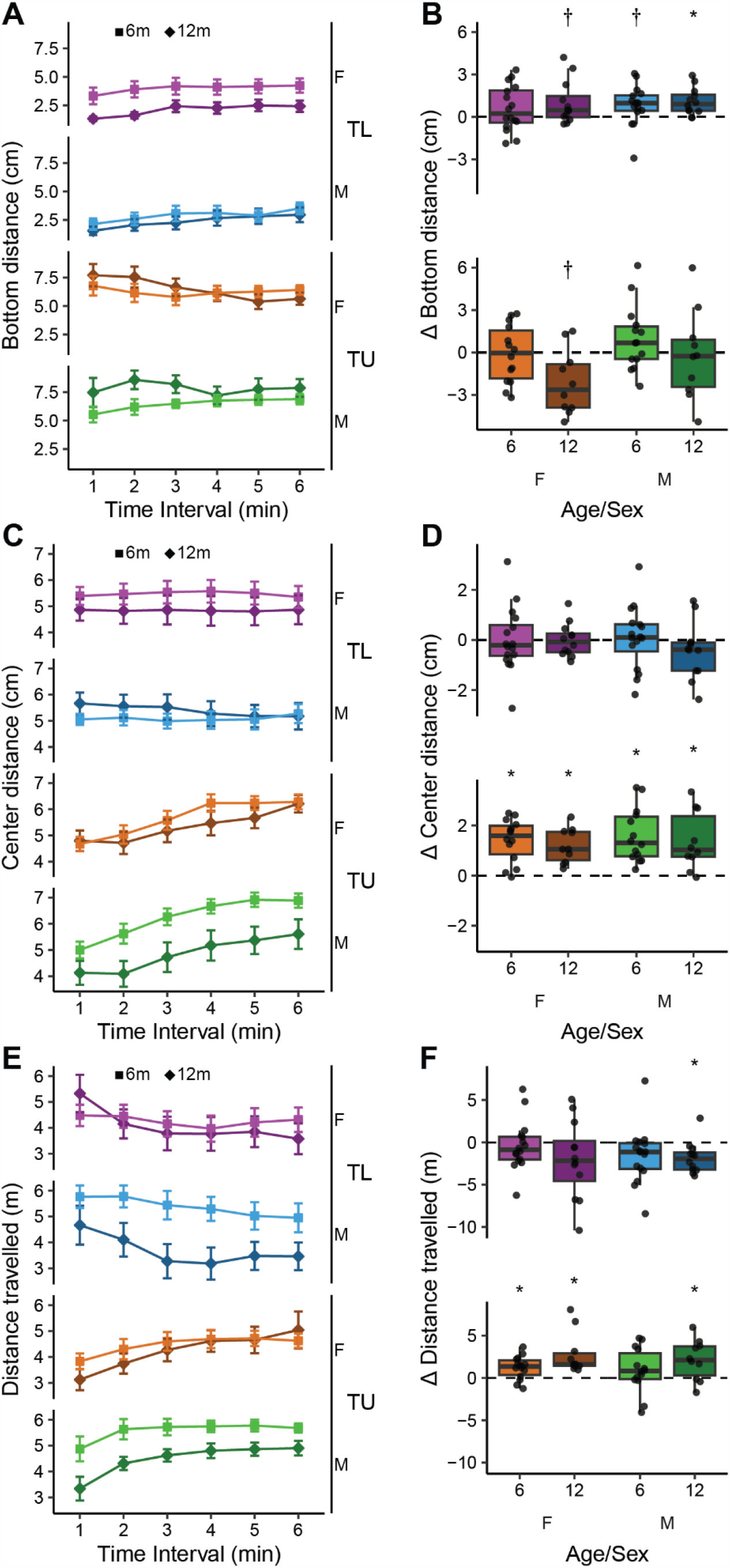
Behavior across time during exploration of the novel tank in young (6 mpf) and old (12 mpf) TL and TU fish of both sexes. Behaviors assessed were (A) minute-by-minute bottom distance, (B) difference in bottom distance during the last two minutes compared to the first two minutes, (C) minute-by-minute center distance, (D) difference in center distance during the last two minutes compared to the first two minutes, (E) minute-by-minute distance travelled, (F) difference in distance travelled during the last two minutes compared to the first two minutes. Boxplot center line is the median, hinges are the interquartile range, and whiskers are hinges ± 1.5 times the interquartile range. *P < 0.05, †P < 0.10 compared to zero using one-sample t-tests with FDR corrections.

For change in distance from the center of the tank over time, TL fish did not appear to habituate whereas TU fish increased center distance over time (Figure 2C). In examining the change in center distance between the first two and last two minutes, in TL fish (Figure 2D, top) there were no effects of age (P = 0.40), sex (P = 0.66), or an interaction (P = 0.42). Furthermore, none of the TL fish showed changes in center distance over time (P’s > 0.84). Likewise, in TU fish (Figure 2D, bottom), there were also no effects of age (P = 0.44), sex (P = 0.50) or an interaction (P = 0.99). However, in contrast to the TL fish, in all groups of the TU fish there were large effects of increased center distance over time (P’s < 0.01, d’s > 1).

Locomotor activity has also been found to change during exposure to a novel environment (Gerlai et al., 2006; Levin et al., 2007). An initial look at the minute-by-minute data (Figure 2E) suggests that TL and TU fish differ in that TL fish decrease distance travelled over time whereas TU fish increase distance travelled. In TL fish, there were no effects of age (P = 0.29), sex (P = 0.52) or an interaction (P = 0.44) on the change in distance travelled over time (Figure 2F, top). Only older male fish significantly decreased over time (P = 0.024, d = 0.98; all other P’s > 0.15). In TU fish, there was a medium sized effect of age (P = 0.039, η^2^ = 0.092) on the change in distance travelled (Figure 2F, bottom) such that older fish increased their distance travelled between the beginning and end of the session more than younger fish. There was no effect of sex (P = 0.49) or an interaction (P = 0.73). When comparing distance travelled in the first two minutes to the last two minutes, there were strong effects of increased locomotor activity over time in young female (P = 0.012, d = 0.87), older female (P = 0.012, d = 1.14) and older male fish (P = 0.027, d = 0.89). There was no change in young male fish (P = 0.20).

### The effect of age on anxiety-related predator-avoidance behaviors during a second exposure to the novel tank

We next sought to determine if the effects of age we saw during the initial exposure to the novel tank persisted during a second exposure 24 hours later. For bottom distance in TL fish (Figure 3A, top) there was no longer a main effect of age (P = 0.20). There was also no effect of sex (P = 0.29), but there was a strong trend towards a medium-sized interaction between age and sex (P = 0.055, η^2^ = 0.069) where older female fish had lower bottom distance than their younger counterparts with the opposite trend in males. Following up the interaction with post-hoc tests within sex indicated a trend towards a large effect in older versus younger TL females (P = 0.088, d = 0.80) and no effect in males (P = 0.58). Thus, the higher anxiety-like behavior in older females, but not males, weakly persisted from day 1 to day 2 in TL fish. In TU fish there were no effects of age (P = 0.51) or sex (P = 0.46) on bottom distance on the second day, although there was a trend towards a medium-sized interaction between age and sex (P = 0.081, η^2^ = 0.067; Figure 3A, bottom) mirroring what was observed in TL fish. However, posthoc tests within sex found no differences between young and old fish in either females (P = 0.43) or males (P = 0.28).

**Figure 3.**
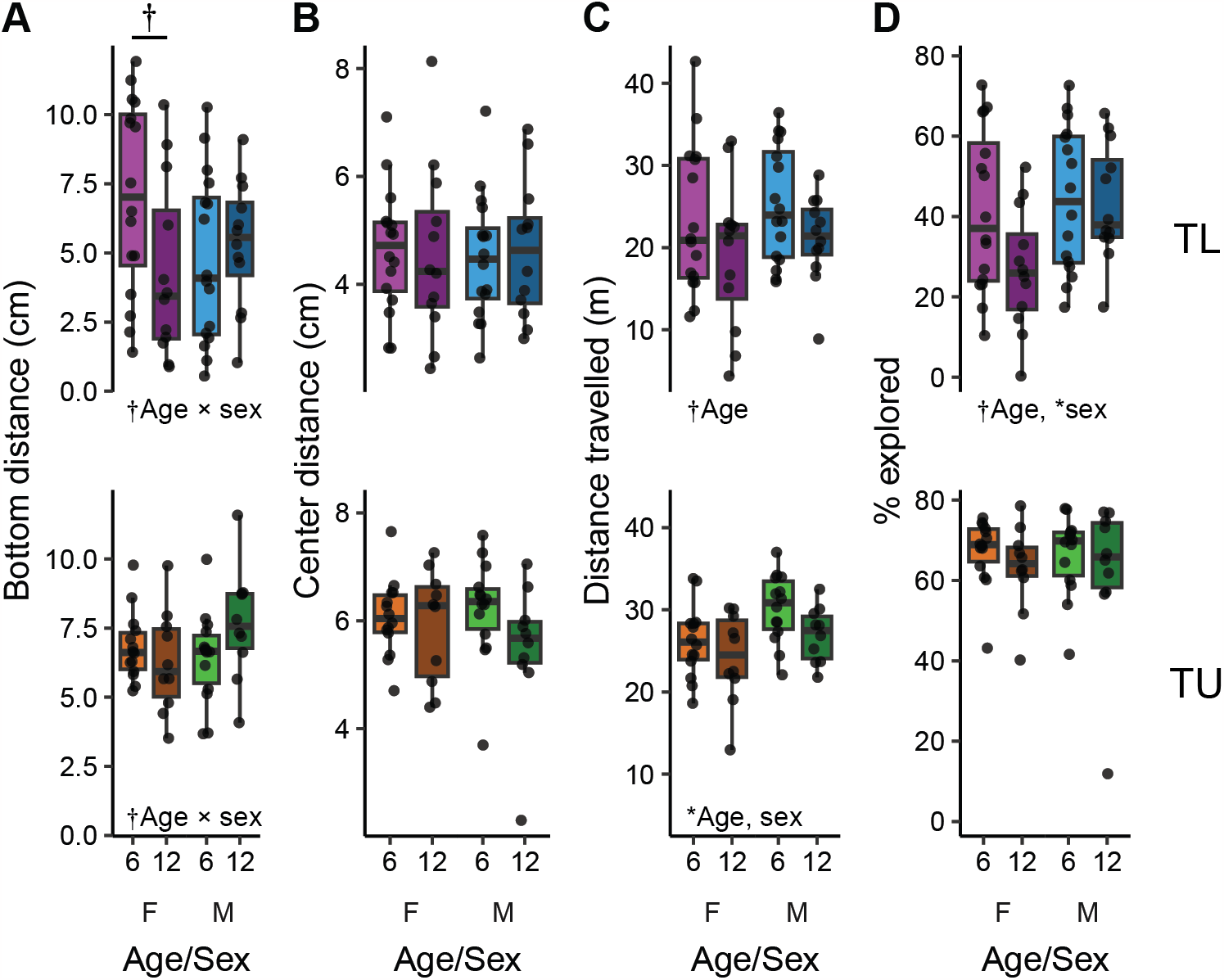
Overall behavior during a second exposure to the novel tank in young (6 mpf) and old (12 mpf) TL (top) and TU (bottom) fish of both sexes. Behaviors assessed were (A) bottom distance, (B) center distance, (C) distance travelled and (D) percent of the tank explored. Boxplot center line is the median, hinges are the interquartile range, and whiskers are hinges ± 1.5 times the interquartile range. *P < 0.05, †P < 0.10 from 2 × 2 (age × sex) ANOVAs.

In TL fish on day 2, center distance (Figure 3B, top) was not influenced by age (P = 0.92), sex (P = 0.90) nor was there an interaction between age and sex (P = 0.75). In TUs, there was no longer an effect of age on center distance on day 2 (P = 0.13), nor was there an effect of sex (P = 0.76) or an interaction (P = 0.33).

### The effects of age on locomotion and exploration during a second exposure to the tank

In TL fish, as on the first day, there was a trend towards a medium-sized effect of age on distance travelled (P = 0.054, η^2^ = 0.068) where older fish swam less than younger fish. There was no effect of sex (P = 0.34) or an interaction (P = 1; Figure 3C, top). The effects of age on locomotor activity in TU fish also persisted to the second day (Figure 3C, bottom): there was a medium-sized effect of age (P = 0.049, η^2^ =0.074) such that older fish swam less than younger fish. There was also a large effect of sex (P = 0.011, η^2^ =0.14) with male fish swimming further than female fish. There was no interaction between age and sex (P = 0.64). Overall, the effects of age and sex on locomotor activity observed on day 2 were largely consistent with what was seen on day 1 suggesting these behavioral differences are particularly stable.

The exploration of an environment may also change upon repeated exposure due to increased familiarity. During the second exposure to the tank, in TL fish there were trends towards small effects of age (P = 0.10, η^2^ =0.046) and sex (P = 0.068, η^2^ = 0.058) with no interaction (P = 0.16). Older TL fish explored less of the tank than younger fish and males explored more of the tank than females (Figure 3D, top). These findings largely mirrored the effects seen on day 1 for age, but now males appeared to explore more of the tank than females. On the second day in TU fish there were no effects of age (P = 0.24), sex (P = 0.81) or an interaction (P = 0.98). Taken together, as we saw on day 1, the influence of age on exploration of the novel tank was more prominent in TLs albeit with smaller effect sizes than observed on day 1.

### The effects of age on long-term habituation

To determine if fish exhibited long-term habituation, we calculated how their behavior changed on day 2 compared to day 1 (Figure 4). In TL fish, we found that all groups of animals increased their bottom distance on the second day (P’s < 0.026, d’s: 0.74 - 1.23; Figure 4A, top). There were no effects of age (P = 0.88), sex (P = 0.64) or an interaction (P = 0.24) on the change in distance from bottom. In contrast, bottom distance in TU fish did not habituate on the second day (P’s > 0.47; Figure 4A, bottom), and there were no effects of age (P = 0.22), sex (P = 0.43) or an interaction (P = 0.46). This lack of an effect in TUs may represent a ceiling effect as they had higher baseline bottom distance on day 1 (Figure 1A, bottom).

**Figure 4.**
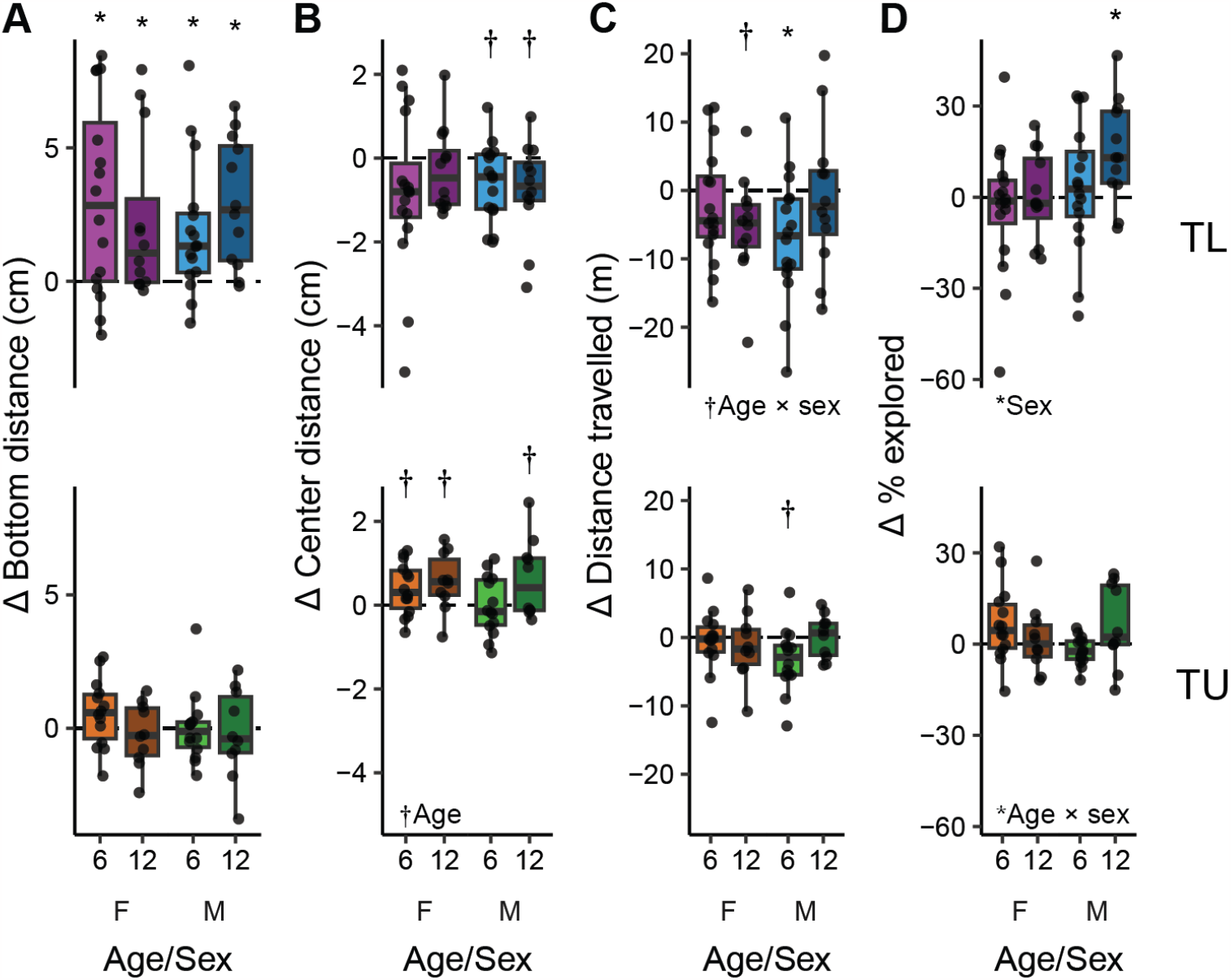
Change in behavior between the first and second days of the NTT in young (6 mpf) and old (12 mpf) TL (top) and TU (bottom) fish of both sexes. Behaviors assessed were changes in (A) bottom distance, (B) center distance, (C) distance travelled and (D) percent of the tank explored. Boxplot center line is the median, hinges are the interquartile range, and whiskers are hinges ± 1.5 times the interquartile range. *P < 0.05, †P < 0.10 from 2 × 2 (age × sex) ANOVAs (below figure) or from one-sample t-tests compared to zero with FDR corrections (above individual boxplots).

For long-term habituation of center distance, all groups of TL fish had medians below zero, decreasing their center distance (Figure 4B, top), although only males of both ages approached statistical significance (P’s = 0.084; d’s = 0.64 and 0.66 for young and old males, respectively; females: P’s > 0.13). There were no effects of age (P = 0.59), sex (P = 0.86) or an interaction (P = 0.33) on the change in center distance in TL fish. In contrast, most TU fish increased their center distance on day 2 (Figure 4B, bottom). There were trends towards medium-sized effects in females (P’s = 0.070, d = 0.63 & 0.80 for young and older females, respectively) and older males (P = 0.084, d = 0.67), but not younger males (P = 0.94). There was also a trend towards a medium-sized effect of age on the change in center distance (P = 0.057, η^2^ = 0.73) such that older animals had a greater increase in center distance than younger animals. There were no effects of sex (P = 0.43) or an interaction (P = 0.28).

The median locomotor activity in TL fish also trended towards a decrease on the second day (Figure 4C, top), although there was only a significant decrease in young males (P = 0.031, d = 0.77) and a trend in older females (P = 0.058, d = 0.72) with no effect in younger females (P = 0.38) or older males (P = 0.74). When comparing the groups to each other, there were no main effects of age (P = 0.55) or sex (P = 0.73), but there was a trend towards a medium sized interaction of age and sex (P = 0.070, η^2^ = 0.062) where, compared to their younger counterparts, older female fish decreased, and older male fish increased, locomotor activity from day 1 to day 2. However, follow-up post-hoc tests within sex found no differences in females (P = 0.32) or males (P = 0.28). Like the TLs, young TU males had a trend towards a decrease in locomotor activity on the second day (P = 0.085, d = 0.70) with no change in any other groups (P’s > 0.86; Figure 4C, bottom). The change in locomotion in TUs was not affected by age (P = 0.31), sex (P = 0.63) and there was no interaction (P = 0.13).

The percentage of the tank explored on the second day changed little in both TL and TU fish (Figure 4D; P’s > 0.24) except for older male TL’s (P = 0.042, d = 0.89). In TL fish there was a trend towards a medium-sized effect of sex on the change in percent explored (P = 0.055, η^2^ =0.069) where males, irrespective of age, increased their percent explored compared to females. There were no effects of age (P = 0.12) or an interaction (P = 0.45). In TU fish, there was no effect of age (P = 0.58) or sex (P = 0.30). There was a medium-sized interaction (P = 0.047, η^2^ =0.87) where young female and older male TUs increased their exploration compared to older females and younger males, however, follow-up post-hoc tests within sex found no differences in females (P = 0.35) or males (P = 0.19).

## Discussion

The behavior of an organism arises through an interaction between environmental conditions and biological factors like genetics, sex, and age. Understanding how these factors interact to influence behavior represents a major challenge in behavioral neurobiology that has important implications for improving human health. Here, we found behavioral changes during exploration of a novel tank in zebrafish that differed in age by only six-months (6 mpf versus 12 mpf). The most consistent behavioral change was a decrease in locomotor activity in older animals. The effects of age on predator avoidance behaviors related to anxiety were more complex as they were sensitive to genetic background and prior exposure to the tank. There were no clear effects of aging on either short-term or long-term habituation.

This is the first study, to our knowledge, to identify changes in predator avoidance behaviors due to aging in zebrafish. We found that the effect of age differed based on strain: older TL fish had increased bottom dwelling (i.e., elevated anxiety-like behavior) whereas older TU fish had a decrease in thigmotaxis (i.e., decreased anxiety-like behavior, but see discussion below on challenges for interpreting thigmotaxis in zebrafish). This complexity in how genetics interacts with age in anxiety-related behaviors has also been found in other model organisms, such as mice, where age is associated with both increases and decreases in anxiety, depending on the specific measure or test, (Shoji et al., 2016), and genetic background (Griebel et al., 2000; Milner and Crabbe, 2008). These findings in animals mirror some of the complexity observed in humans, where genetic contributions to fear and anxiety vary across anxiety phenotypes (Shimada-Sugimoto et al., 2015). Thus, the present study suggests that age-related changes in the behavior of zebrafish can be used to model some aspects of human behavior.

We found that older TU fish had a decrease in thigmotaxis (i.e., spent more time closer to the center of the tank). Thigmotaxis in zebrafish has typically been interpreted as predator avoidance and thus anxiety-related based on its similarity to behavior in rodents (Champagne et al., 2010; Kalueff et al., 2013; Maximino et al., 2010). However, recent findings call this interpretation into question. For example, bottom dwelling is better established as a predator avoidance behavior related to anxiety (Gerlai, 2010; Kalueff et al., 2013; Levin and Cerutti, 2009; Maximino et al., 2010), and when measured concurrently with thigmotaxis, these two parameters do not consistently correlate with each other (Rajput et al., 2022). Furthermore, several studies have found that animals with high levels of freezing or immobility, a well-defined index of fear and anxiety (Blaser et al., 2010; Maximino et al., 2010), spend more time in the center of the tank than the periphery (Wong et al., 2012; Ziv et al., 2013). We, and others, have also found thigmotaxis to increase over time in a novel environment in some fish strains (Fig. 2C; Champagne et al., 2010; Rajput et al., 2022; Shams et al., 2015), which is the opposite of what would be expected as animals habituate. Finally, the anxiolytic and anxiogenic effects of drugs often have the expected effects on bottom dwelling without altering thigmotaxis (Ochocki and Kenney, 2023; Rosa et al., 2018). Taken together, this suggests that the interpretation of thigmotaxis in zebrafish is not straightforward. The present study suggests the interesting possibility that whether thigmotaxis is indicative of predator-avoidance and anxiety may be related to genetic background: TU fish display several changes in thigmotaxis (e.g., short-term habituation [Fig. 2C], long-term habituation [Fig 4B], and the effects of age [Fig. 1B]) whereas effects in TL fish are largely confined to bottom dwelling. When TL fish do have changes in thigmotaxis (i.e., long-term habituation, Fig. 4B) it is reduced instead of increased like it is in TU fish. However, the idea that genetic background may influence thigmotaxis in zebrafish requires further study.

The most consistent effect of age we observed was a decrease in locomotion in older fish. This effect was present in both strains and both days that animals were placed in the tank and is in line with prior work (Gilbert et al., 2014; Philpott et al., 2012; Yu et al., 2006). Indeed, age related decreases in locomotion have been observed in a wide range of species such as fruit flies (Grotewiel et al., 2005; Leffelaar and Grigliatti, 1983), mice (Shoji et al., 2016), and humans (Iosa et al., 2014; Moore, 1975; Vandervoort, 2002). This makes decreased locomotor activity a particularly consistent aging biomarker that can facilitate comparative work (Marck et al., 2017). In zebrafish, age-related decreases in locomotor activity are associated with reduced endurance and swim performance, likely due to sarcopenia (i.e., the loss of muscle mass and strength) (Daya et al., 2020; Gilbert et al., 2014). In humans, sarcopenia is also thought to be a key driver of reduced locomotor activity due to aging (Cruz-Jentoft et al., 2014; Doherty, 2003). Given the overlap in the structure and development of skeletal muscle in zebrafish and mammals (Daya et al., 2020; Keenan and Currie, 2019), our findings lend further support to the use of zebrafish as a model system for studying the effects of aging on muscle function and performance.

We did not observe any clear effects of age on either short-term or long-term habituation memory in our study. This is in contrast to prior work that found older zebrafish had deficits in associative learning (Ruhl et al., 2015; Yu et al., 2006). This discrepancy may be due to the use of different ages. Prior work used animals ranging in age from 1 to 3 years of age whereas in the present study animals ranged in age from 6 to 12 months. It may be the case that the age difference needs to be larger to see changes in learning. Alternatively, it may be that associative learning is more sensitive to the effects of age in zebrafish than the non-associative habituation learning examined in the present study.

As zebrafish grow in prominence as a model organism in behavioral neurobiology (Gerlai, 2023; Kenney, 2020) it is critical to understand how biological factors influence their behavior. This is essential for not only identifying what aspects of human behavior can be modeled, but also for deciding what age, sex, and strain of animal may be best suited to address a specific question. The findings of the present study, that only a modest difference in age can result in several behavioral changes, suggests zebrafish may prove to be a useful model organism for studying aging and that care should be taken when choosing zebrafish strains when studying the effects of age.

## Methods

### Subjects

Animals were female or male TL or TU zebrafish either 6 months post-fertilization (mpf; 24-32 weeks of age) or 12 mpf (52-58 weeks of age). The number of fish per group are as follows with the same number of male and female fish per group: TLs, 6mpf: n’s = 16; TLs, 12mpf: n’s = 12, TU, 6mpf: n’s = 14, TU, 12mpf: n’s = 10. All animals were bred and raised at Wayne State University and were within two generations from fish obtained from the Zebrafish International Resource Center at the University of Oregon. Animals were housed on high density racks using standard conditions (27.5 ± 0.5 °C, 500 ± 10 μS, pH of 7.4 ± 0.2) and a 14:10 light:dark cycle (lights on at 8:00AM). Feeding was twice daily with a dry feed (Gemma 300, Skretting, Westbrook, ME, USA) in the morning and brine shrimp (*Artemia salina*, Brine Shrimp Direct, Ogden, UT, USA) in the afternoon.

Fish were sexed using three secondary sex characteristics: color, shape, and presence of pectoral fin tubercles in males (McMillan et al., 2015). Sex was confirmed after behavioral procedures: fish were euthanized and dissected to determine the presence or absence of eggs. All procedures were approved by the Wayne State University Institutional Animal Care and Use Committee.

### Novel tank test

The novel tank test was performed in five-sided tanks (15 × 15 × 15 cm) made from frosted acrylic (TAP Plastics, Stockton, CA, USA) and filled with 2.5 L of fish facility water (to a height of 12 cm). Tanks were open from above and placed in an enclosure of white plasticore to prevent the influence of external stimuli and to diffuse light. Videos were recorded using D435 Intel RealSense™ depth-sensing cameras (Intel, Santa Clara, CA, USA) connected to Linux workstations. Cameras were mounted 20 cm above the tanks. Fish were tracked using DeepLabCut (Mathis et al., 2018). We generated three-dimensional swim traces and extracted four behavioral parameters (distance from bottom, distance from center, distance travelled, and percent of tank explored) using custom written Python code as previously described (Rajput et al., 2022).

At least one week prior to behavioral testing fish were dual housed as male/female pairs. Two-liter tanks were divided in half using a transparent barrier and pairs of fish were placed in each section of the tank. One hour prior to testing in the novel tank, fish were removed from the racks and allowed to habituate in the procedure room. Fish were then placed individually into the novel tank for six minutes. Water in the tank was changed between animals. Following testing, fish remained in the procedure room for an hour prior to being returned to their racks.

### Statistical analysis

Analysis was done using R version 4.3.0 (R Core Team, 2016) and graphed using ggplot2 (Wickham, 2015). Data were analyzed for significance using 2 × 2 (age × sex) ANOVAs as appropriate. Interactions were followed up with independent samples t-tests within sex and corrected for multiple comparisons using the false discover rate (Benjamini and Hochberg, 1995). When assessing how behavior changed over time, we used one-sample t-tests (μ = 0) and corrected for multiple comparisons as above. Effect sizes are reported as η^2^ for ANOVAs and Cohen’s d for t-tests and interpreted as small (0.01 < η^2^ < 0.06; 0.2 < d < 0.5), medium (0.06 ≤ η^2^ < 0.14; 0.5 ≤ d < 0.8) or large (η^2^ ≥ 0.14; d ≥ 0.8) based on Cohen (1988).

## Acknowledgements

This work was funded by the National Institutes of Health (R35GM142566) to J.W.K.

